# ABA-IMPORTING TRANSPORTER 1.1 activity in the endosperm of tomato seeds restrains germination under salinity stress

**DOI:** 10.1101/2022.02.28.482276

**Authors:** Hagai Shohat, Hadar Cheriker, Amir Cohen, David Weiss

**Author notes:** Author for correspondence: David Weiss,.

## Abstract

The plant hormone abscisic acid (ABA) has a central role in the regulation of seed maturation and dormancy. ABA also restrains germination under abiotic-stress conditions. Here we show in tomato (*Solanum lycopersicum*) that the ABA importer *ABA-IMPORTING TRANSPORTER 1*.*1* (AIT1.1) has a role in radicle emergence following imbibition. *AIT1*.*1* expression was upregulated during imbibition, and CRISPR/Cas9-derived *ait1*.*1* mutant seeds exhibited faster radicle emergence, increased germination and partial resistance to ABA. *AIT1*.*1* was highly expressed in the endosperm, but not in the embryo, and *ait1*.*1* isolated embryos did not show resistance to ABA. On the other hand, loss of AIT1.1 activity promoted the expression of endosperm-weakening related genes, and seed-coat scarification eliminated the promoting effect of *ait1*.*1* on radicle emergence. Therefore, we propose that imbibition-induced *AIT1*.*1* expression in the endosperm, mediates ABA-uptake into endosperm cells to restrain endosperm weakening. While salinity conditions strongly inhibited the wild-type (WT) M82 seed germination, it had much weaker effect on *ait1*.*1* germination. We suggest that this AIT1.1 function was evolved to inhibit germination under unfavorable conditions, such as salinity. Unlike other ABA mutants, *ait1*.*1* exhibited normal seed longevity, and therefore, the *ait1*.*1* allele may have a potential for improving seed germination in crops.

## Introduction

The timing of seed germination is important for seedling establishment (Donohue et al., 2010), and due to its irreversible nature, this process is highly regulated. Germination is regulated by environmental and internal cues including light, temperature, water availability and hormones. The plant hormones gibberellin (GA) and abscisic acid (ABA) promotes and inhibits germination, respectively (Nonogaki, 2014).

ABA accumulates during seed maturation and induces seed dormancy (Finch-Savage and Leubner-Metzger, 2006). ABA-induced seed dormancy is orchestrated by dormancy-related genes such as *ABSCISIC ACID INSENSITIVE 3 (ABI3), FUSCA 3 (FUS3), DELAY OF GERMINATION1 (DOG1)* and *LEAFY COTYLEDON 1 (LEC1)* and *LEC2* (Holdsworth et al., 2008). The output of these regulatory machinery promotes storage compound accumulation, inhibits viviparous germination, reduces metabolic activity, and leads to the acquisition of desiccation tolerance, enabling embryo survival under long periods of dry environment (Graeber et al., 2012).

During the early stages of seed imbibition, dormancy is released due to ABA catabolism, which precedes the accumulation of GA (Toyomasu et al., 1998; Ogawa et al., 2003; Nambara and Marion-Poll, 2005; Raghavendra et al., 2010). These increase GA to ABA ratio and promote seed germination (Piskurewicz et al., 2008; Liu and Hou, 2018, Shu et al., 2018). In dicots seeds, high GA to ABA ratio induces the accumulation of cell-wall-loosening enzymes, leading to endosperm and seed coat (testa) weakening (Muller et al., 2006). GA also promotes radicle elongation, and the growing radicle ruptures the weakened endosperm and testa, leading to radicle emergence and germination (Steinbrecher and Leubner-Metzger, 2018). Unfavorable conditions, such as salinity, inhibit ABA catabolism, reduce GA to ABA ratio and inhibit endosperm weakening, embryo growth and germination (Shu et al., 2017; Ruggiero et al., 2019; Lai et al., 2020).

ABA activity is regulated at the level of metabolism and signaling, but also by its transport to target tissues and cells (Anfang and Shani, 2021). The first characterized ABA transporters were the ATP-BINDING CASSETTE (ABC) transporters ABCG25 (Kuromori et al., 2010) and ABCG40 (Kang et al., 2010). ABCG25 is expressed mainly in vascular tissues and functions as an exporter. Loss of ABCG25 resulted in reduced germination. ABCG40, on the other hand, is an ABA importer that is expressed in guard cells. The *abcg40* mutant closed its stomata slower than the wild type (WT) in response to ABA, exhibited reduced tolerance to drought and increased germination on ABA. The integrated activity of the ABCG transporters controls seed germination in Arabidopsis and Medicago (Kang et al., 2015; Pawela et al., 2019). The ABA transporter Detoxification Efflux Carriers (DTX)/Multidrug and Toxic Compound Extrusion (MATE) DTX50 is upregulated by ABA and expressed mainly in guard cells and vascular tissue (Zhang et al., 2014). The *dtx50* mutant plants exhibited hypersensitivity to ABA during seed germination and increased tolerance to drought. Additional ABA transporter from the NITRATE TRANSPORTER1 (NRT1)/PTR TRANSPORTER FAMILY (NPF) named ABA-IMPORTING TRANSPORTER 1 (AIT1) or NRT1.2 was characterized in Arabidopsis (Kanno, et al., 2012). *AIT1* is expressed mainly in vascular tissues of inflorescence stem, and the *ait1* mutant exhibits increased transpiration due to larger stomatal aperture and a partial resistance to ABA (Shimizu et al., 2021). A tomato *AIT1* (*AIT1*.*1*) homologue was recently characterized (Shohat et al., 2020). CRISPR/Cas9-derived tomato *ait1*.*1* mutant exhibits increased transpiration and reduced stomatal closure in response to ABA.

Tomato seeds exhibit primary non-deep physiological dormancy, and can be stored dry for years (Hilhorst and Downie, 1995; Baskin and Baskin, 2004). The tomato embryo is enclosed in a rigid endosperm and testa which act as a physical barrier to restrict radicle emergence (Yan et al., 2014). During seed imbibition, the endosperm at the radicle tip region (i.e. micropylar endosperm) is weakened due to GA-induced cell-wall-loosening enzyme activity, enabling radicle emergence (Nonogaki et al., 2000). Here, we show that the tomato *AIT1*.*1* is upregulated in the endosperm during imbibition. The *ait1*.*1* mutant seeds exhibited improved germination under normal and salinity conditions. Our results suggest that AIT1.1 acts in the endosperm during imbibition to mediate ABA-suppression of endosperm weakening and germination under normal and salinity conditions.

## Results

### *AIT1*.*1* is upregulated during tomato seed imbibition to restrain germination

ABA accumulates during seed maturation and promotes seed dormancy, whereas following imbibition, it is catabolized to enable germination (Braybrook and Harada, 2008). To study the possible role of the ABA transporter AIT1.1 in seed maturation, dormancy and/or germination, we first examined its expression in WT M82 seeds during different developmental and physiological stages. We collected seeds from green, color break and red ripped tomato fruits, dry seeds one day, 2 weeks and 4 weeks after extraction from red ripen fruits, and at different times during seed imbibition (0-96h). *AIT1*.*1* expression was low during seed maturation and slightly upregulated in dry seeds (Fig. 1A). It was strongly upregulated during imbibition (up to 72h), and following radicle emergence (96h), its expression was downregulated. These results imply that AIT1.1 has a role during seed imbibition, perhaps to restrain germination.

**Fig. 1.**
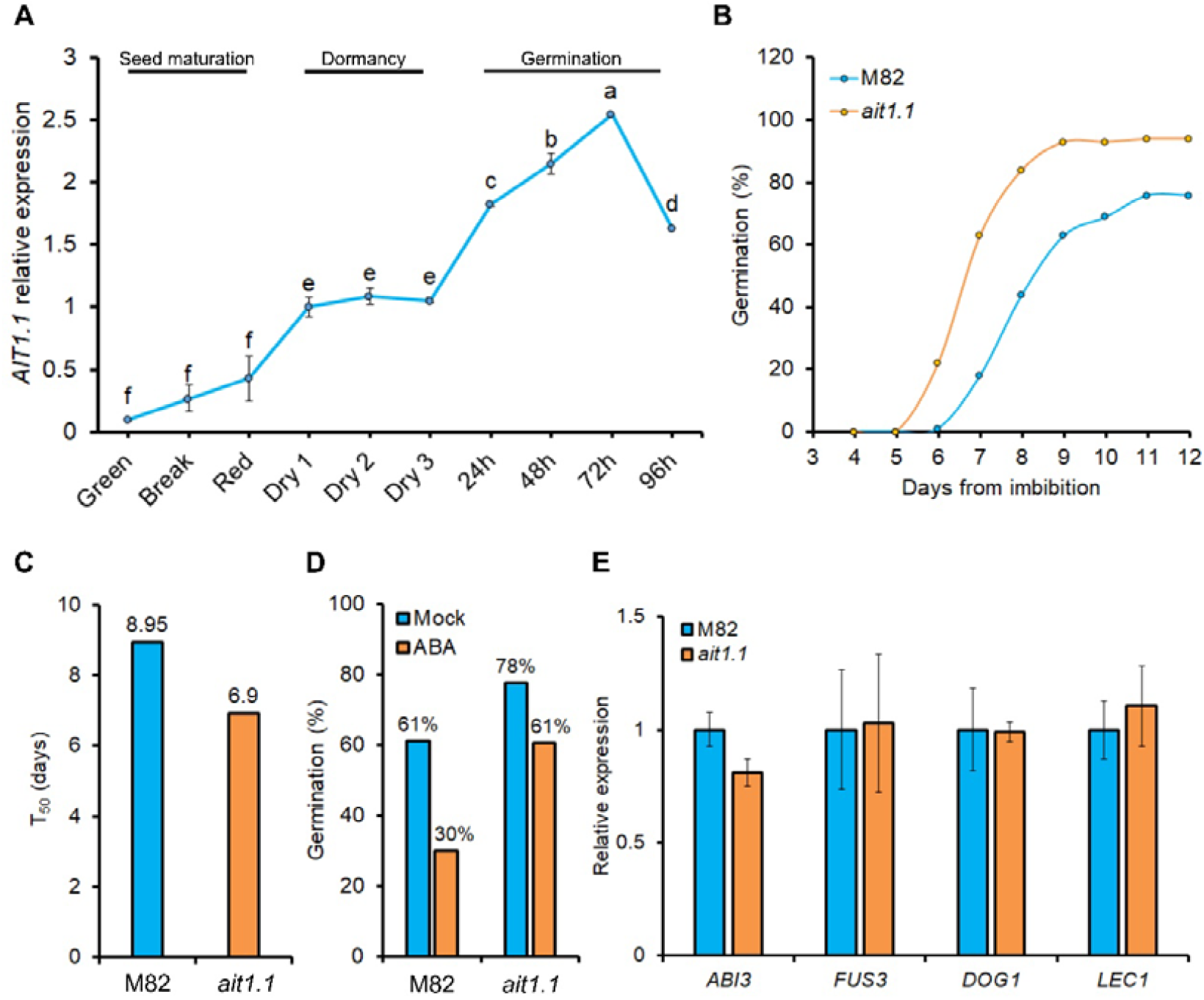
*AIT1*.*1* is upregulated during seed imbibition and restrains germination. (A) Relative expression of *AIT1*.*1* in seeds taken from green, color break and red ripped tomato fruits, as well as, from fresh dry seeds for one day (dry 1), 2 weeks (dry 2) or 4 weeks (dry 3) after extraction from red ripen fruits, and imbibed seeds (0-96h imbibition after 4 weeks of dry storage). (B) Germination of M82 and *ait1*.*1* seeds in soil. (C) Germination rate (T_50_) of M82 and *ait1*.*1*. (D) Percentages of germinated seeds after 10 days in petri dishes containing MS medium with (or without) 1µM ABA. (E) Relative expression of central regulators of seed dormancy genes *ABI3, FUS3, DOG1* and *LEC1* in M82 and *ait1*.*1* dry seeds stored for 1 month. Values in A and E are means of 4 replicates, each contains 10 seeds ±SE. Values in B, C and D are percentages or means of 100 (B,C) or 50 (D) seeds. Small letters (A) represent significant differences between respective treatments by Student’s *t* test (*P* < *0*.*05*).

We next examined whether the loss of AIT1.1 activity in *ait1*.*1* CRISPR/Cas9-derived mutant (Shohat et al., 2020) affects germination. The mutant seeds started to germinate 6 days after sowing in the soil, and reached 93% germination after 8 days, whereas M82 seeds started to germinate after 7 days, reaching 78% germination after 11 days (Fig. 1B). Germination rate can be determined by the time (days) takes to reach 50% germination (T_50_; Coolbear et al., 1984). T_50_ of *ait1*.*1* was 6.9, whereas that of M82 was 8.95 (Fig. 1C). In a germination assay on MS media containing ABA, *ait1*.*1* seed exhibited partial resistance to the hormone and germinated at higher percentage than M82 (Fig. 1D).

We tested whether *ait1*.*1* also affects dormancy by analyzing the transcriptional profile of the central regulators of seed dormancy *ABI3, FUS3, DOG1* and *LEC1*. Loss of AIT1.1 activity had no effect on the expression of these genes (Fig. 1E). It was shown before that reduced ABA accumulation or activity decreases seed desiccation tolerance and therefore longevity (Ooms et al., 1993). We tested the germination of M82 and *ait1*.*1* seeds, that were stored for one or 15 months. The loss of *AIT1*.*1* did not affect the longevity of the seeds, and they exhibited normal germination after the long storage, suggesting that *AIT1*.*1* has no role in seed desiccation tolerance (Fig. S1). Seeds of the ABA-deficient *sitiens* mutant (*sit*, Groot and Karssen, 1992) on the other hand, exhibited reduced germination after long storage (15 months, Fig. S2). Taken together, the results suggest that *AIT1*.*1* does not affect dormancy or desiccation tolerance, but probably imbibition-related processes.

Tomato has four *AIT1* genes (Shohat et al., 2020) and we examined their expression pattern during imbibition to uncover possible functional redundancy. While *AIT1*.*1* and its closest homolog *AIT1*.*2* were upregulated during imbibition, *AIT1*.*3* and *AIT1*.*4* were downregulated (Fig. 2A), suggesting that different *AIT1* genes in tomato have different role in seeds. We next generated CRISPR/Cas9-derived *ait1*.*2* mutant. The *ait1*.*2* mutant gene had 5bp deletion causing a frame shift and a premature stop codon (Fig. S3). Homozygous mutant plants did not show any clear phenotypic alteration. We performed germination assay and found that *ait1*.*2* seeds exhibit reduced germination compared to M82 (Fig. 2B). These results ruled out the possibility of functional redundancy with *AIT1*.*1*. It is possible that AIT1.2, opposite to AIT1.1, acts as an ABA exporter.

**Fig. 2.**
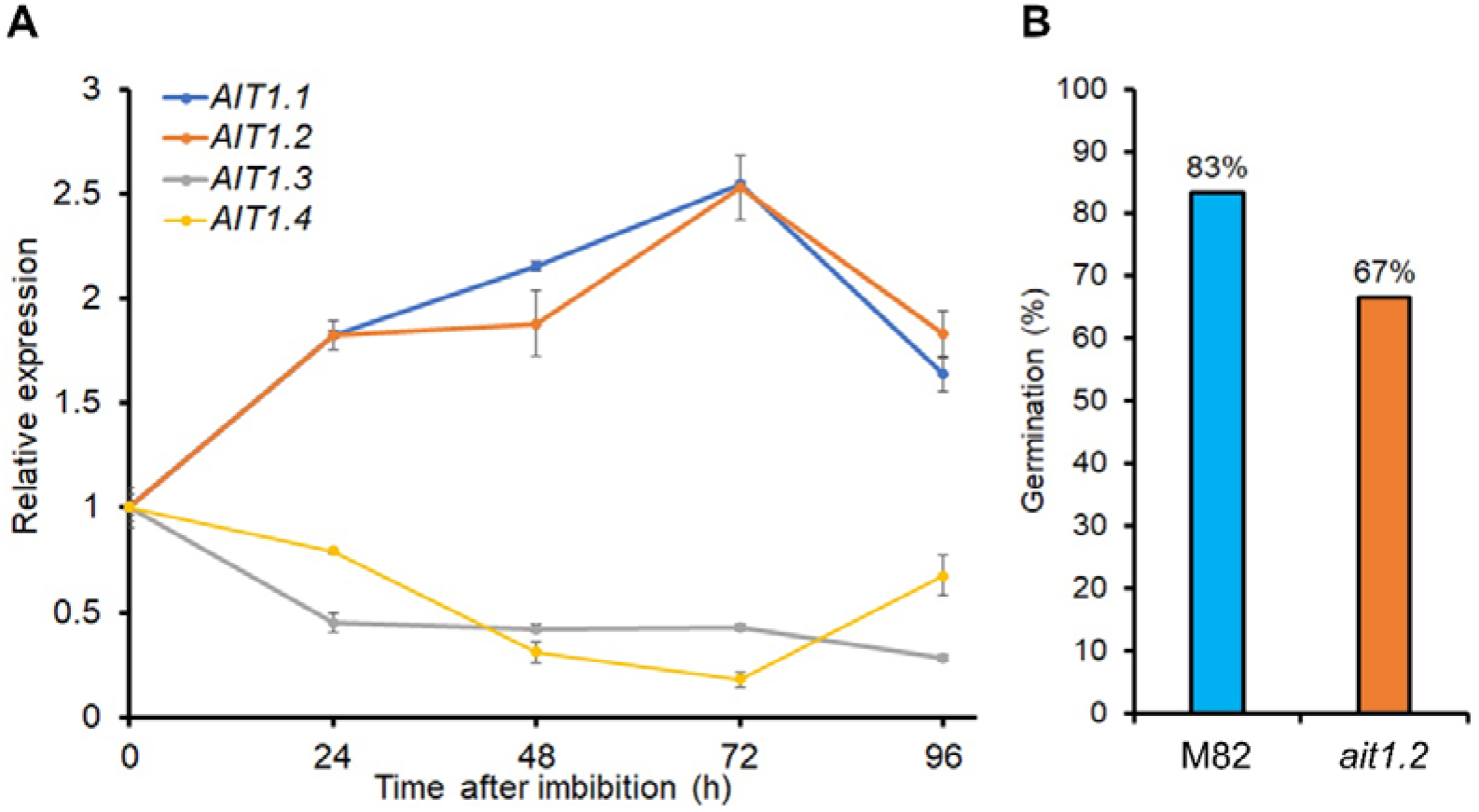
Expression dynamics of the tomato *AIT1* genes during seed imbibition. (A) Relative expression of the tomato *AIT1* genes in seeds during imbibition (0-96h imbibition after 4 weeks of dry storage). Values are means of 4 replicates each contains 10 seeds ±SE. (B) Germination of M82 and *ait1*.*2* seeds in soil after 10 days. Values are percentages of 70 seeds.

### AIT1.1 acts in the endosperm to mediate seed germination

Tomato seed germination is controlled by two parallel processes; embryo growth and endosperm weakening (Steinbrecher and Leubner-Metzger, 2017). We first tested whether AIT1.1 acts in the embryo to mediate seed germination. To this aim, we imbibed M82 and *ait1*.*1* seeds for several hours, then rescued the embryos from the seeds and grew them for 72h on MS medium with or without ABA (Fig. 3A). We analyzed two ABA-inhibited developmental processes; embryo growth and greening. We did not find differences in growth and greening between M82 and *ait1*.*1* embryos on MS with or without ABA (Fig. 3B,C), suggesting that AIT1.1 does not affect germination via its activity in the embryo.

**Fig. 3.**
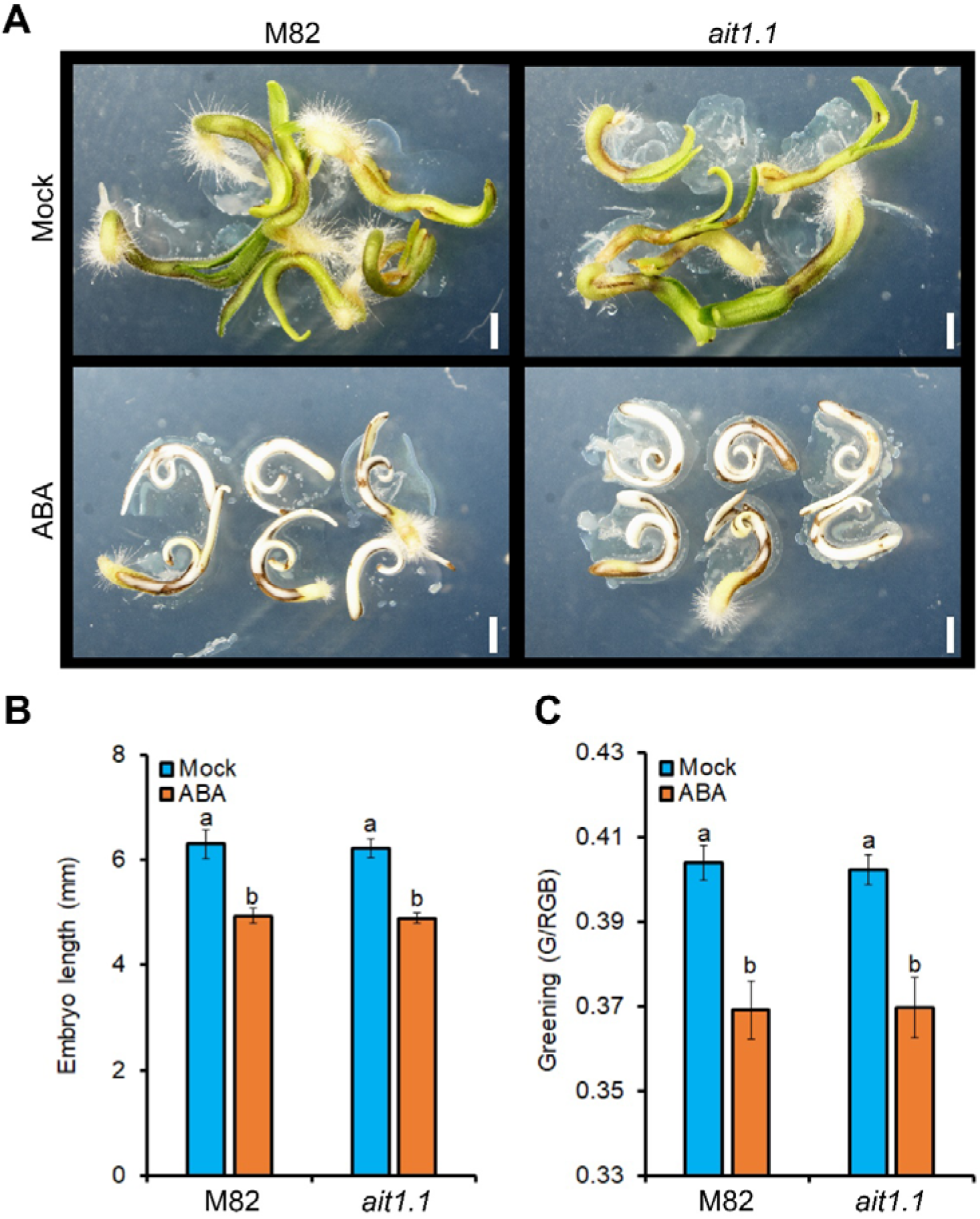
AIT1.1 did not affect embryo growth. (A) Representative isolated embryos grown for 72h on petri dishes containing MS medium with or without 10µM ABA. (B) Length of M82 and *ait1*.*1* isolated embryos grown for 72h on Petri dishes containing MS medium with (or without) 10µM ABA. (C) Greening of M82 and *ait1*.*1* isolated embryos grown for 72h on Petri dishes containing MS medium with (or without) 10µM ABA. Values are means of 10 biological replicates ±SE. Small letters in B and C represent significant differences between respective treatments by Tukey-Kramer HSD test (*P* < *0*.*05)*. Scale bar in A = 1mm.

The Arabidopsis ABCG ABA transporters controls ABA transport from the endosperm to the embryo to regulate embryo growth and germination (Kang et al., 2015). To test whether AIT1.1 has a similar role in tomato, we have used the Seed Coat Bedding Assay (SCBA, Lee et al., 2010) that tests the inhibition of embryo growth by endosperm-synthesized ABA. We found that isolated M82 embryos taken from imbibed seeds, developed faster on MS medium compared to embryos that were placed on a layer of embryo-less seed coats and endosperms (Fig. 4A,B). However, we did not find differences in embryo growth and greening between M82 and *ait1*.*1* in reciprocal SCBA assays (Fig. 4C, Fig. S4), suggesting that the increased germination in *ait1*.*1* is probably not a result of inhibition of ABA transport from the endosperm to the embryo.

**Fig. 4.**
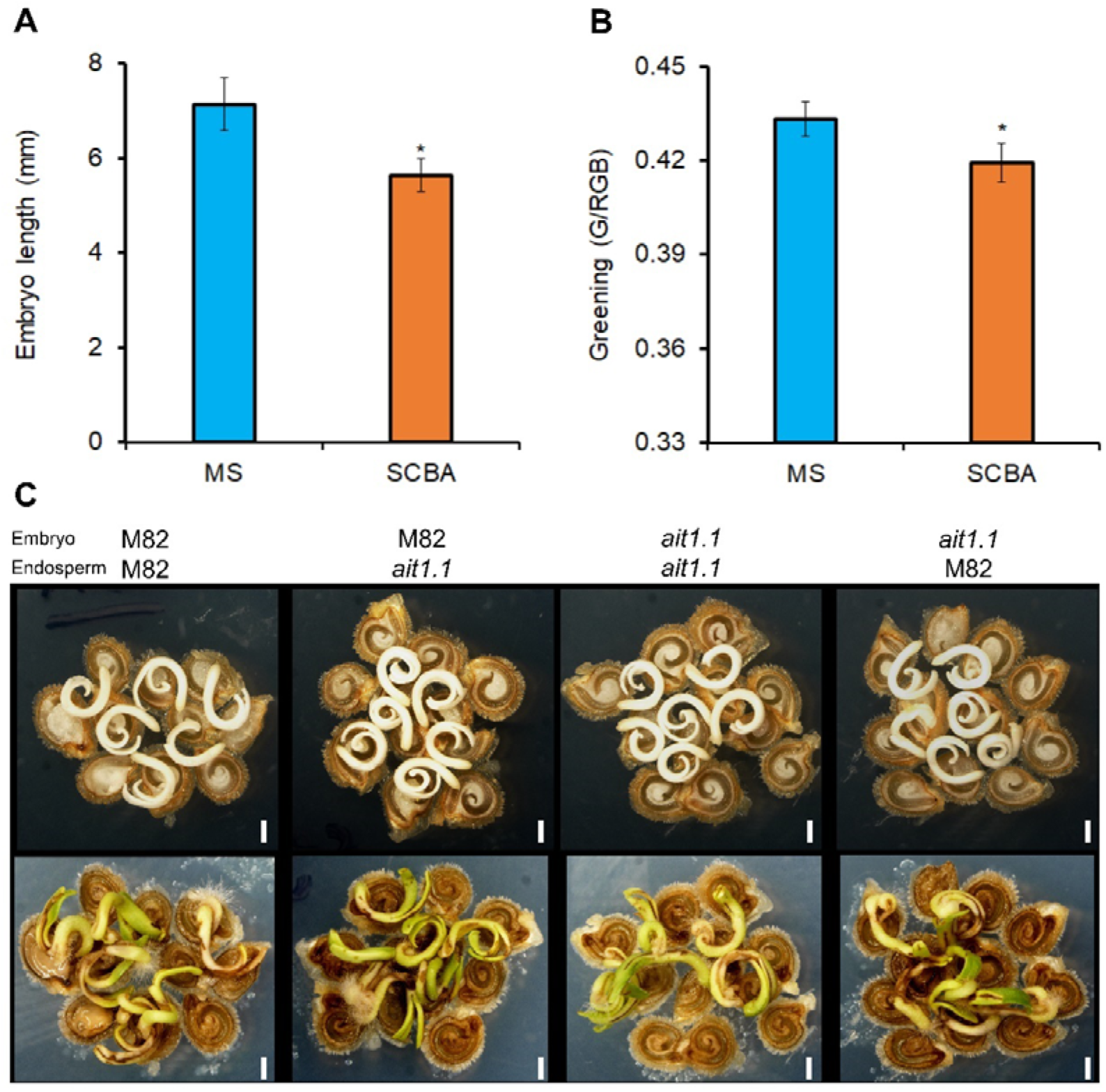
AIT1.1 did not affect endosperm inhibition of embryo growth. (A) Length of M82 isolated embryos grown for 72h on MS medium or on a layer of endosperm containing seed coat (seed coat bedding assay-SCBA). (B) Greening of M82 isolated embryos grown for 72h on MS medium or on a layer of endosperm containing seed coat (SCBA). (C) Representative M82 and *ait1*.*1* reciprocal SCBA assays grown for 72h on Petri dishes containing MS medium. Values are means of 5 (A,B) or 10 (C) biological replicates ± SE. Stars in A and B represent significant differences between respective treatments by Student’s t test (*P* < *0*.*05*). Scale bar = 1mm.

*AIT1*.*1* expression was significantly higher in the endosperm compared to the embryo (Fig. 5A). Moreover, GUS activity in transgenic *AIT1*.*1*_*pro*_:GUS (Shohat et al., 2020) imbibed seeds was significantly higher in the endosperm compared to the embryo (Fig. 5B). We therefore tested whether the increased germination of *ait1*.*1* seeds is a result of increased endosperm weakening. We first scarified the seed at the micropylar region to eliminate the inhibitory effect of the endosperm and the testa on radicle emergence. Following scarification, we did not find differences between M82 and *ait1*.*1* in the time of radicle emergence or in post-emergence primary root growth with or without ABA (Fig. 5C, Fig. S5). We then analyzed the expression of endosperm-weakening-related genes *EXPANSIN2* (*EXPA2*), β*1,3-Glucanase and ENDO-BETA-MANNASE 2* (*MAN2*) during imbibition (Leubner-Metzger et al., 1995; Martinez-Andujar et al., 2012; Graeber et al., 2014). These genes exhibited higher expression in *ait1*.*1* endosperm, suggesting higher endosperm weakening activity in the mutant (Fig. 5D).

**Fig. 5.**
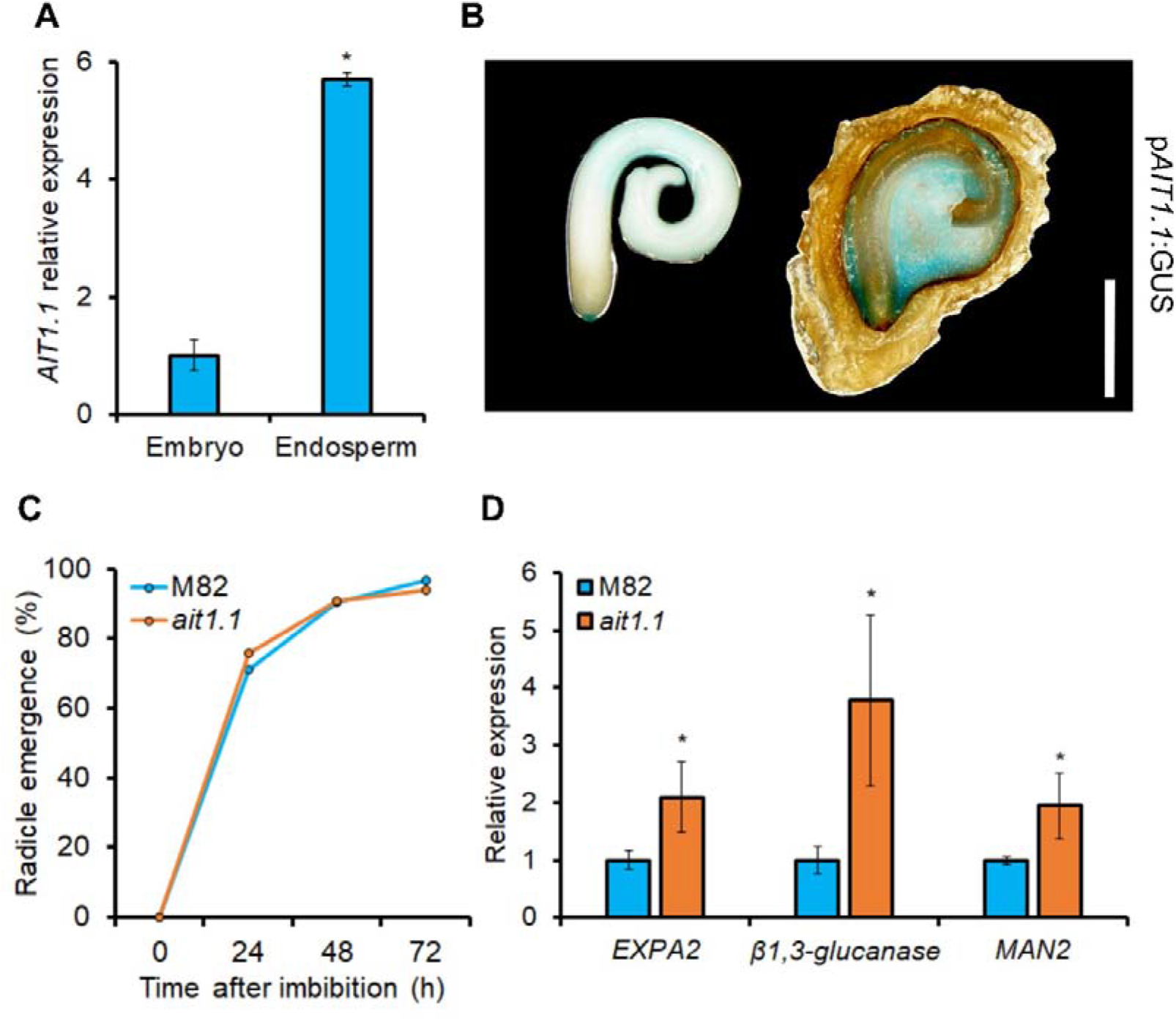
AIT1.1 acts in the endosperm to restrain radicle emergence. (A) Relative expression of *AIT1*.*1* in isolated embryos or endosperms after 48h of imbibition. (B) Representative *pAIT1*.*1:GUS* transgenic embryo and embryo-less endosperm showing GUS staining after 48h of imbibition. (C) Percentages of radicle emergence in M82 and *ait1*.*1* seeds following scarification. (B) Relative expression of endosperm weakening marker genes *EXPA2*, □*1,3-glucanase* and *MAN2* in endosperms of M82 and *ait1*.*1* after 48h of imbibition. Values in A and D are means of 4 replicates each contains 10 seeds ± SE. Values in C are percentages from 30 seeds. Stars in A and D represent significant differences between respective treatments by Student’s *t* test (*P* < *0*.*05*). Scale bar = 1mm.

### *AIT1*.*1* restrained germination under salt stress conditions

Salinity conditions suppress germination via the inhibition of ABA catabolism (Kong et al., 2017; Xia et al., 2019). We tested if AIT1.1 has a role in the inhibition of germination imposed by salinity. M82 and *ait1*.*1* seeds were germinated on 0, 25, 50, 75 and 100 mM NaCl. In the presence of low NaCl concentrations (0 and 25 mM), *ait1*.*1* seeds were germinated slightly better than M82 seeds. However, in the presence of higher NaCl concentration (50 mM) M82 seeds exhibited 20% and *ait1*.*1* exhibited 60% germination (Fig. 6). We did not find any effect of salt or ABA on *AIT1*.*1* expression in imbibed seeds (Fig. S6), suggesting that although *AIT1*.*1* plays a role in the inhibition of germination under salt stress, it is not regulated by salinity or ABA at the transcriptional level.

**Fig. 6.**
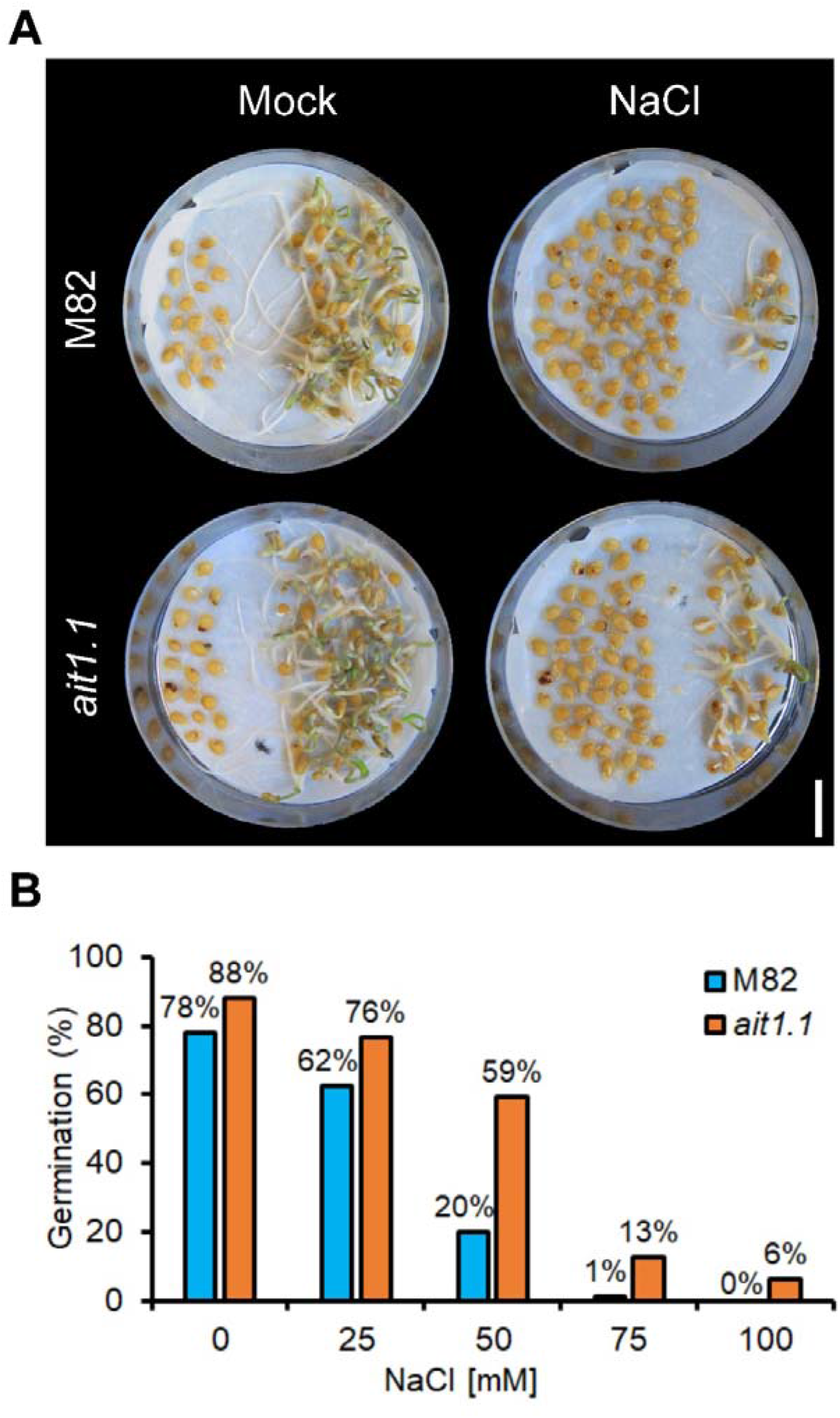
Loss of *AIT1*.*1* promoted germination under salinity conditions. (A) Representative M82 and *ait1*.*1* seeds placed for 5 days on Petri dishes with or without 50mM NaCl. (B) Percentages of germinated seeds placed for 7 days on Petri dishes with or without different concentrations of NaCl. Values are percentages of 80 seeds. Scale bar in A = 1cm.

## Discussion

ABA levels decrease in seeds during imbibition to enable germination (Nambara and Marion-Poll, 2005). However, under salinity-stress conditions, ABA levels remain high and inhibit germination (Kong et al., 2017). All these are eliminated in ABA-deficient mutants that are able to germinate on salt-containing media (Groot and Karssen, 1992; Léon-Kloosterziel et al., 1996). Here we show in tomato, that the ABA importer *AIT1*.*1* acts in the endosperm to mediate the inhibitory effect of ABA on germination under salinity conditions.

Although ABA is the central regulator of seed maturation, dormancy and desiccation tolerance, the expression of *AIT1*.*1* was relatively low during these stages, and was significantly upregulated during imbibition, when ABA level decreases (Braybrook and Harada, 2008). The Arabidopsis *AIT1* homolog exhibits similar expression pattern (http://bar.utoronto.ca/efp/cgi-bin/efpWeb.cgi). Loss of *AIT1*.*1* did not affect the expression of the major dormancy-related genes, but increased the expression of genes related to endosperm weakening. These suggest that *AIT1*.*1* restrains seed germination, but has no role in ABA-mediated seed maturation or dormancy. In line with this, the loss of *AIT1*.*1* had no effect on seed longevity and therefore, it probably has no role in seed desiccation tolerance.

Tomato has four *AIT1* genes (Shohat et al., 2020). While *AIT1*.*1* and *AIT1*.*2* were upregulated during imbibition, *AIT1*.*3* and *AIT1*.*4* were downregulated. This opposite expression dynamics of *AIT1* genes implies that they might have roles in different processes, e.g., *AIT1*.*1* and *AIT1*.*2* in germination and *AIT1*.*3/AIT1*.*4* in maturation, dormancy or desiccation tolerance. Although *AIT1*.*1* and *AIT1*.*2* have similar expression pattern they seem to have opposite role; while the loss of *AIT1*.*1* improves germination, the loss of *AIT1*.*2* inhibits germination. We previously showed that AIT1.1 acts as an ABA importer. It is possible that AIT1.2 is an ABA exporter; in Arabidopsis, the ABA transporter ABCG40 is an importer, whereas its homolog ABCG31 is an exporter (Kang et al., 2015)

The process of seed germination includes two major paralleled processes; embryo growth and endosperm weakening, both are inhibited by ABA (Lee et al., 2012; Belin et al., 2019). We found that the loss of *AIT1*.*1* had no effect on the growth of isolated embryos or on radicle emergence upon scarification. It was previously suggested that ABA is synthesized in the endosperm and transported to the embryo to restrict its growth during germination (Kang et al., 2015). However, our SCBA assays do not support such a role for AIT1.1 in tomato seeds. All together these results suggest that *AIT1*.*1* does not affect embryo growth. *AIT1*.*1* was predominantly expressed in the endosperm and the expression of endosperm weakening marker genes *EXPA2*, [*1,3-glucanase* and *MAN2* was upregulated in *ait1*.*1* endosperm. These suggest that AIT1.1 mediates the inhibitory effect of ABA on endosperm weakening and radicle emergence.

Although the loss of *AIT1*.*1* promoted germination under normal conditions, its effect was much more prominent under salt stress. It is possible that the main role of AIT1.1 in seeds is to transport ABA during imbibition under unfavorable conditions to inhibit germination. Although ABA and salinity did not affect the expression of *AIT1*.*1* in seeds, it is possible that the high ABA levels under salinity conditions, post-translationally regulate AIT1.1 activity to increase ABA transport into endosperm cells. It was recently shown that ABA activates a kinase that phosphorylates the Arabidopsis AIT1 to enhance ABA transport and responses (Zhang et al., 2021). Taken together we propose that imbibition under salinity conditions increases ABA levels in tomato seeds, and AIT1.1 facilitates ABA-uptake into the micropylar endosperm to block the effects of GA on endosperm weakening, radicle emergence and germination.

Plants have adopted mechanisms to reduce germination under unfavorable conditions. These promote the survival of a population under changing environment (Gioria et al., 2018; Vishal and Kumar, 2018). However, this evolutionary advantage, is an undesired trait for agricultural needs (Finch-Savage and Bassel, 2016; Fabrissin et al., 2021). The *ait1*.*1* mutant exhibited increased and uniform germination, but had no effect on seed longevity. Moreover, in field experiment we did not find any effect of the mutation on fruit quality, size and total yield (Fig. S7). These results suggest that this mutation can be harnessed to promote uniform and rapid germination without affecting seed longevity and fruit yield in commercial tomato varieties.

## Materials and Methods

### Plant materials, growth conditions and hormone treatments

Tomato (*Solanum lycopersicum*) plants in M82 background (*sp/sp*) were used throughout this study. *ait1*.*1* (Shohat et al., 2020), *ait1*.*2, sit* (Nir et al., 2017) and the transgenic line *pAIT1*.*1:GUS* (Shohat et al., 2020) were backcrossed to or generated in M82 background. Plants were grown in a growth room set to a photoperiod of 12/12-h night/day, light intensity of 150 μmol m^-2^ s^-1^ and 25°C and irrigated to saturation. The seeds were harvested from ripped fruits and treated with 1% sodium hypochlorite followed by 1% Na_3_PO_4_ 12H_2_O, and incubated with 10% Sucrose over-night in 37°C. Seeds were stored dry at room temperature. (±)-ABA dissolved in dimethyl sulfoxide (DMSO, Sigma-Aldrich, St. Louis, USA) and NaCl dissolved in DDW.

### CRISPR/Cas9 mutagenesis, tomato transformation and selection of mutant alleles

Four single-guide RNAs (sgRNAs, Supplemental Table 2) were designed to target *AIT1*.*2* gene, using the CRISPR-P tool (http://cbi.hzau.edu.cn/crispr). Vectors were assembled using the Golden Gate cloning system, as described by Weber et al. (2011). Final binary vectors pAGM4723 was introduced into *A. tumefaciens* strain GV3101 by electroporation. The constructs were transferred into M82 cotyledons using transformation and regeneration methods described by McCormick (1991). Kanamycin-resistant T0 plants were grown and independent transgenic lines were selected and self-pollinated to generate homozygous transgenic lines. The genomic DNA of each plant was extracted and genotyped by PCR for the presence of the Cas9 construct. The CRISPR/Cas9-positive lines were further genotyped for mutations using a forward primer to the upstream sequence of the sgRNA1 target and a reverse primer to the downstream of the sgRNA2 target sequence. *AIT1*.*2* was sequenced in all mutant lines. Homozygous line was identified and selected for further analysis. The Cas9 construct was segregated out by crosses to M82.

### Germination assays

Seeds were germinated in soil or petri dishes contained Murashige & Skoog (MS) medium (Duchefa, Haarlem, NL) or wet Whatman filter paper (GE Healthcare, Amersham, UK) in a growth room set to a photoperiod of 12/12-h night/day, with a light intensity of 150 µmol m^-2^ s^-1^, and at a temperature of 25°C. In soil, germination was scored when the hypocotyl emerged from the soil, and in petri dishes when the radicle pierced the seed coat.

### RNA extraction and cDNA synthesis

Total RNA extracted by RNeasy Plant Mini Kit (Qiagen). For synthesis of cDNA, SuperScript II reverse transcriptase (18064014; Invitrogen, Waltham, MA, USA) and up to 3 mg of total RNA were used, according to the manufacturer’s instructions.

### RT-qPCR analysis

RT-qPCR analysis was performed using an Absolute Blue qPCR SYBR Green ROX Mix (AB-4162/B) kit (Thermo Fisher Scientific, Waltham, MA 15 USA). Reactions were performed using a Rotor-Gene 6000 cycler (Corbett Research, Sydney, Australia). A standard curve was obtained using dilutions of the cDNA sample. The expression was quantified using Corbett Research Rotor-Gene software. Three independent technical repeats were performed for each sample. Relative expression was calculated by dividing the expression level of the examined gene by that of *SlACTIN*. Gene to *ACTIN* ratio was then averaged. All primers sequences are presented in Supplemental Table 1. The values of mock and/or M82 WT treatments were set to 1.

### Embryo and radicle length measurements

Embryos and radicles images were captured using a Nikon SMZ1270 stereo microscope equipped with a Nikon DS-Ri2 camera and NIS-ELEMENT software (Nikon Instruments, Melville, USA). Images were than analyzed to determine length using ImageJ software segmented line tool (http://rsb.info.nih.gov/ij/).

### Embryo greening analysis

Embryos greening was analyzed using ImageJ software histogram feature. Greening was determined by calculating the green pixels intensity relative to RGB intensity. The minimum greening value which calculated from white screen was 0.34, and the maximum value which calculated from 4 days old embryo cotyledons was 0.54.

### Embryos isolation and Seed Coat Bedding Assay (SCBA)

To isolate embryos, tomato seeds were imbibed for several hours in water, and then the seeds were dissected into embryos and endosperms by using a surgical blade and very fine tweezers under a binocular microscope (Olympus, Waltham, MA, USA). The SCBAs were performed according to Lee et al., (2010). Briefly, isolated embryos were placed on a layer of embryo-less endosperms that laid on a MS medium. Embryos growth and greening were analyzed after 72h as described above (See material and Methods).

### GUS staining

Histochemical detection of GUS activity was performed using 5-bromo-4-chloro-3-indolyl-b-D-glucuronide as described in Donnelly et al., (1999). Samples were photographed under a Nikon SMZ1270 stereo microscope equipped with a Nikon DS-Ri2 camera and NIS-ELEMENT software.

### Statistical analyses

All assays were conducted with three or more biological replicates and analyzed using JMP software (SAS Institute, Cary, NC). Means comparison was conducted using ANOVA with post-hoc Tukey-Kramer HSD (For multiple comparisons) and Student’s t tests (For one comparison) (P<0.05).

## Accession Numbers

Sequence data from this article can be found in the Sol Genomics Network (https://solgenomics.net/) under the following accession numbers: *ACTIN, Solyc11g005330; AIT1*.*1, Solyc05g006990; AIT1*.*2, Solyc05g007000; AIT1*.*3, Solyc04g005790; AIT1*.*4, Solyc03g113250*; *ABI3, Solyc06g083600; FUS3, Solyc02g094460; DOG1, Solyc03g006120; LEC1, Solyc04g015060;* β*1-3-glucanase, Solyc01g060010; EXPA2, Solyc09g018020; MAN2, Solyc06g064520*.

## Acknowledgments

This research was supported by the Israel Science Foundation (617/20) to DW.

## Author contributions

HS and DW designed the research plan; HS, HC and AC performed the research; HS and AC analyzed data; HS and DW wrote the paper.

## Data availability

All data can be found in the manuscript and in the supporting information.

## References

Anfang M, Shani E. 2021. Transport mechanisms of plant hormones. Current Opinion in Plant Biology 63, 102055.

Baskin JM, Baskin CC. 2004. A classification system for seed dormancy. Seed Science Research 14, 1–16.

Belin C, Megies C, Hauserova E, Lopez-Molina L. 2009. Abscisic acid represses growth of the Arabidopsis embryonic axis after germination by enhancing auxin signaling. The Plant Cell 21, 2253–2268.

Braybrook SA, Harada JJ. 2008. LECs go crazy in embryo development. Trends in Plant Science 13, 624–630.

Coolbear P, Francis A, Grierson D. 1984. The effect of low temperature pre-sowing treatment on the germination performance and membrane integrity of artificially aged tomato seeds. Journal of Experimental Botany 35, 1609–1617.

Donnelly PM, Bonetta D, Tsukaya H, Dengler RE, Dengler NG. 1999. Cell cycling and cell enlargement in developing leaves of Arabidopsis. Developmental Biology 215, 407–419.

Donohue K, de Casas RR, Burghardt L, Kovach K, Willis CG. 2015. Germination, postgermination adaptation, and species ecological ranges. Annual Review of Ecology, Evolution, and Systematics 41, 293–319.

Fabrissin I, Sano N, Seo M, North HM. 2021. Ageing beautifully: can the benefits of seed priming be separated from a reduced lifespan trade-off? Journal of Experimental Botany 72, 2312–2333.

Finch-Savage WE, Bassel GW. 2016. Seed vigour and crop establishment: extending performance beyond adaptation. Journal of Experimental Botany 67, 567–591.

Finch-Savage WE, Leubner-Metzger G. 2006. Seed dormancy and the control of germination 171, 501–523.

Gioria M, Pysek P, Osborne BA. 2018. Timing is everything: does early and late germination favor invasions by herbaceous alien plants? Journal of Plant Ecology 11, 4–16.

Graeber K, Linkies A, Steinbrecher T, et al. 2014. DELAY OF GERMINATION 1 mediates a conserved coat-dormancy mechanism for the temperature-and gibberellin-dependent control of seed germination. Proceedings of the National Academy of Sciences, USA 111, E3571–E3580.

Graeber K, Nakabayashi K, Miatton E, Leubner-Metzger G, Soppe WJJ. 2012. Molecular mechanisms of seed dormancy. Plant, Cell & Environment 35, 1769–1786.

Groot SPC, Karssen CM. 1992. Dormancy and germination of abscisic acid-deficient tomato seeds. Plant Physiology 99, 952–958.

Hilhorst HWM, Downie B. 1995. Primary dormancy in tomato (Lycopersicon esculentum cv. Moneymaker): studies with the sitiens mutant. Journal of Experimental Botany 47, 89–97.

Holdsworth MJ, Bentsink L, Soppe WJJ. 2008. Molecular networks regulating Arabidopsis seed maturation, after-ripening, dormancy and germination. New Phytologist 179, 33–54.

Kang J, Hwang JU, Lee M, Kim YY, Assmann S, Martinoia E, Lee Y. 2010. PDR-type ABC transporter mediates cellular uptake of the phytohormone abscisic acid. Proceedings of the National Academy of Sciences, USA 107, 2355–2360.

Kang J, Yim S, Choi H, Kim A, Lee KP, Lopez-Molina L, Martinoia E, Lee Y. 2015. Abscisic acid transporters cooperate to control seed germination. Nature Communications 6, 8113.

Kanno Y, Hanada A, Chiba Y, Ichikawa T, Nakazawa M, Matsui M, Koshiba T, Kamiya Y, Seo M. 2012. Identification of an abscisic acid transporter by functional screening using the receptor complex as a sensor. Proceedings of the National Academy of Sciences, USA 109, 9653–9658.

Kong X, Luo Z, Zhang Y, Li W, Dong H. 2017. Soaking in H2O2 regulates ABA biosynthesis and GA catabolism in germinating cotton seeds under salt stress. Acta Physiologiae Plantarum 39, 2.

Kuromori T, Miyaji T, Yabuuchi H, Shimizu H, Sugimoto E, Kamiya A, Moriyama Y, Shinozaki K. 2010. ABC transporter AtABCG25 is involved in abscisic acid transport and responses. Proceedings of the National Academy of Sciences, USA 107, 2361–2366.

Lai Y, Zhang D, Wang J, Wang J, Ren P, Yao L, Si E, Kong Y, Wang H. 2020. Integrative transcriptomic and proteomic analyses of molecular mechanism responding to salt stress during seed germination in Hulless Barley. International Journal of Molecular Sciences 21, 359.

Lee KP, Piskurewicz U, Tureckova V, Carat S, Chappuis R, Strnad M, Fankhauser C, Lopez-Molina L. 2012. Spatially and genetically distinct control of seed germination by phytochromes A and B. Genes & Development 26, 1984–1996.

Lee KP, Piskurewicz U, Tureckova V, Strnad M, Lopez-Molina L. 2010. A seed coat bedding assay shows that RGL2-dependent release of abscisic acid by the endosperm controls embryo growth in Arabidopsis dormant seeds. Proceedings of the National Academy of Sciences, USA 107, 19108–19113.

Léon-Kloosterziel KM, Gil MA, Ruijs GS, Jacobsen SE, Olszewski NE, Schwartz SH, Zeevaart JA, Koornneef M. 1996. Isolation and characterization of abscisic acid-deficient Arabidopsis mutants at two new loci. The Plant Journal 10, 655–661.

Leubner-Metzger G, Frundt C, Vogeli-Lange R, Meins F. 1995. Class I LJ-1,3-Glucanases in the endosperm of Tobacco during germination. Plant physiology 109, 751–759.

Liu X, Hou X. 2018. Antagonistic regulation of ABA and GA in metabolism and signaling pathways. Frontiers in Plant Science 9, 251.

Martinez-Andujar C, Pluskota WE, Bassel GW, et al. 2013. Mechanisms of hormonal regulation of endosperm cap-specific gene expression in tomato seeds. The Plant Journal 71, 575–586.

McCormick S. 1991. Transformation of tomato with Agrobacterium tumefaciens. Plant Tissue Culture Manual B6, 1–9.

Muller K, Tintelnot S, Leubner-Metzger G. 2006. Endosperm-limited Brassicaceae seed germination: abscisic acid inhibits embryo-induced endosperm weakening of Lepidium sativum (cress) and endosperm rupture of Cress and Arabidopsis thaliana. Plant Cell Physiology 47, 864–877.

Nambara E, Marion-Poll A. 2005 Abscisic acid biosynthesis and catabolism. Annual Review of Plant Biology 56, 165–185.

Nir I, Shohat H, Panizel I, Olszewski N, Aharoni A, Weiss D. 2017. The tomato DELLA protein PROCERA acts in guard cells to promote stomatal closure. The Plant Cell 29, 3186–3197.

Nonogaki H. 2014. Seed dormancy and germination-emerging mechanisms and new hypotheses. Frontiers in Plant Science 5, 1–14.

Nonogaki H, Gee OH, Bradford KJ. 2000. A germination-specific Endo-b-Mannanase gene is expressed in the micropylar endosperm cap of tomato seeds. Plant Physiology 123, 1235–1245.

Ogawa M, Hanada A, Yamauchi Y, Kuwahara A, Kamiya Y, Yamaguchi S. 2003. Gibberellin biosynthesis and response during Arabidopsis seed germination. The Plant Cell 15, 1591–604.

Ooms JJJ, Leon-Kloosterziel KM, Bartels D, Koornneef M, Karssen CM. 1993. Acquisition of desiccation tolerance and longevity in seeds of Arabidopsis thaliana. Plant Physiology 102, 1185–1191.

Pawela A, Banasiak J, Biala W, Martinoia E, Jasinski M. 2019. MtABCG20 is an ABA exporter influencing root morphology and seed germination of Medicago truncatula. The Plant Journal 98, 511–523.

Piskurewicz U, Jikumaru Y, Kinoshita N, Nambara E, Kamiya Y, Lopez-Molina L. 2008. The gibberellic acid signaling repressor RGL2 inhibits Arabidopsis seed germination by stimulating abscisic acid synthesis and ABI5 activity. The Plant Cell 20, 2729–2745.

Raghavendra AS, Gonugunta VK, Christmann A, Grill E. 2010. ABA perception and signaling. Trends in Plant Science 15, 395–401.

Ruggiero A, Landi S, Punzo P, Possenti M, Van Oosten MJ, Costa A, Morelli G, Maggio A, Grillo S, Batelli G. 2019. Salinity and ABA seed responses in pepper: expression and interaction of ABA core signaling components. Frontiers in Plant Science 10, 304.

Shimizu T, Kanno Y, Suzuki H, Watanabe S, Seo, M. 2021. Arabidopsis NPF4.6 and NPF5.1 control leaf stomatal aperture by regulating abscisic acid transport. Genes 12, 885.

Shohat H, Illouz Eliaz N, Kanno Y, Seo M, Weiss D. 2020. The tomato DELLA protein PROCERA promotes abscisic acid responses in guard cells by upregulating an abscisic acid transporter. Plant Physiology 184, 518–528.

Shu K, Qi Y, Chen F, Meng Y, Luo X, Shuai H, Zhou W, Ding J, Du J, Liu J, Yang F, Wang Q, Liu W, Yong T, Wang X, Feng Y, Yang W. 2017. Salt stress represses soybean seed germination by negatively regulating GA biosynthesis while positively mediating ABA biosynthesis. Frontiers in Plant Science 8, 1372.

Shu K, Zhou W, Chen F, Luo X, Yang W. 2018. Abscisic acid and gibberellins antagonistically mediate plant development and abiotic stress responses. Frontiers in Plant Science 9, 416.

Steinbrecher T, Leubner-Metzger G. 2017. The biomechanics of seed germination. Journal of Experimental Botany 68, 765–783.

Steinbrecher T, Leubner-Metzger G. 2018. Tissue and cellular mechanics of seeds. Current Opinion in Genetics & Development 51, 1–10.

Toyomasu T, Kawaide H, Mitsuhashi W, Inoue Y, Kamiya Y. 1998. Phytochrome regulates gibberellin biosynthesis during germination of photoblastic lettuce seeds. Plant Physiology 118, 1517–1523.

Vishal B, Kumar PP. 2018. Regulation of seed germination and abiotic stresses by gibberellins and abscisic acid. Frontiers in Plant Science 9, 838.

Weber E, Engler C, Gruetzner R. Werner S, Marillonnet S. 2011. A modular cloning system for standardized assembly of multigene constructs. PLoS ONE 6, e16765.

Xia K, Liu A, Wang Y, Yang W, Jin Y. 2019. Mechanism of salt-inhibited early seed germination analysed by transcriptomic sequencing. Seed Science Research 29, 73–84.

Yan D, Duermeyer L, Leoveanu C, Nambara E. 2014. The functions of the endosperm during seed germination. Plant Cell Physiology 55, 1521–1533.

Zhang H, Zhu H, Pan Y, Yu Y, Luan S, Li L. 2014. A DTX/MATE-type transporter facilitates abscisic acid efflux and modulates ABA sensitivity and drought tolerance in Arabidopsis. Molecular Plant 7, 1522–1532.

Zhang L, Yu Z, Xu Y, et al. 2021. Regulation of the stability and ABA import activity of NRT1.2/NPF4.6 by CEPR2-mediated phosphorylation in Arabidopsis. Molecular Plant 14, 633–646.

