## Supplemental Data for "ABA-IMPORTING TRANSPORTER 1.1 activity in the endosperm of tomato seeds restrains germination under salinity stress"

**Fig. S1.** Loss of *AIT1.1* did not affect seed longevity.

**Fig. S2.** ABA deficiency reduced seed longevity.

**Fig. S3.** Sequence analysis of *ait1.2* CRISPR mutant allele.

**Fig. S4.** AIT1.1 did not affect endosperm inhibition of embryo growth.

**Fig. S5.** Loss of *AIT1.1* did not affect post-emergence radicle elongation.

**Fig. S6.** Salt or ABA treatments did not affect *AIT1.1* expression in imbibed seeds.

**Fig. S7.** Loss of *AIT1.1* did not affect yield production.

**Table S1.** Primers used in this study.

**Table S2.** gRNAs used in this study.

**Supplemental Figures**

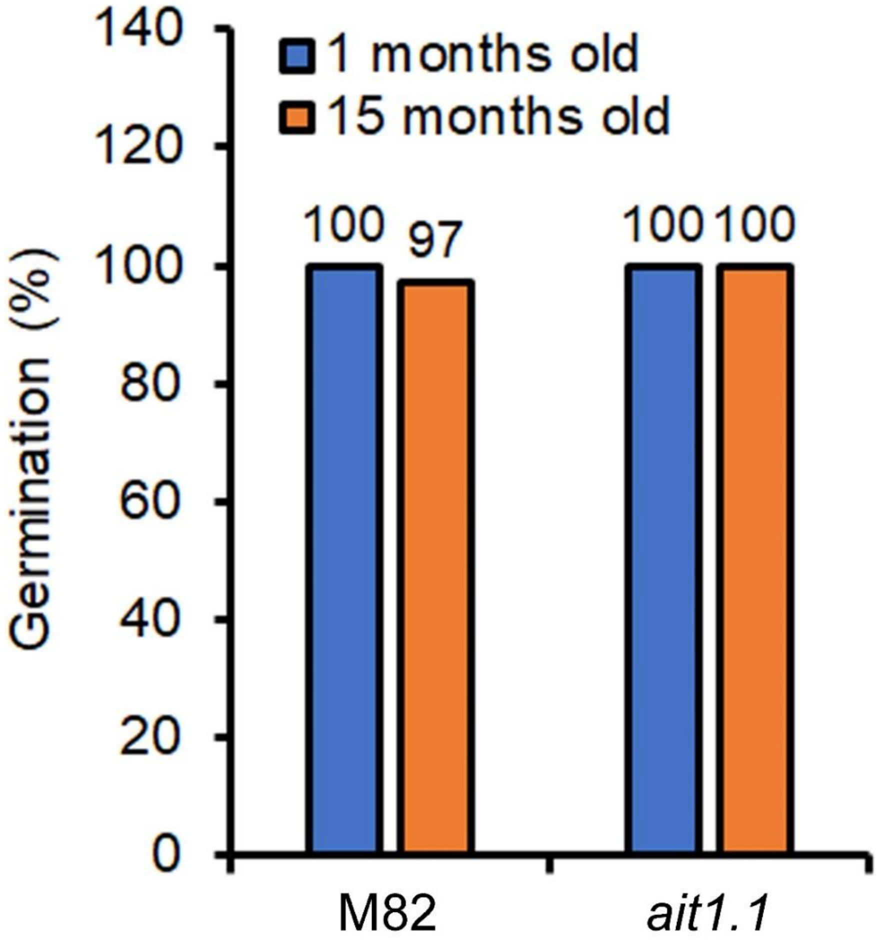


**Fig. S1** Loss of *AIT1.1* did not affect seed longevity. Germination of M82 and *ait1.1* seeds stored for 1 or 15 months. Values are percentages from 50 seeds.

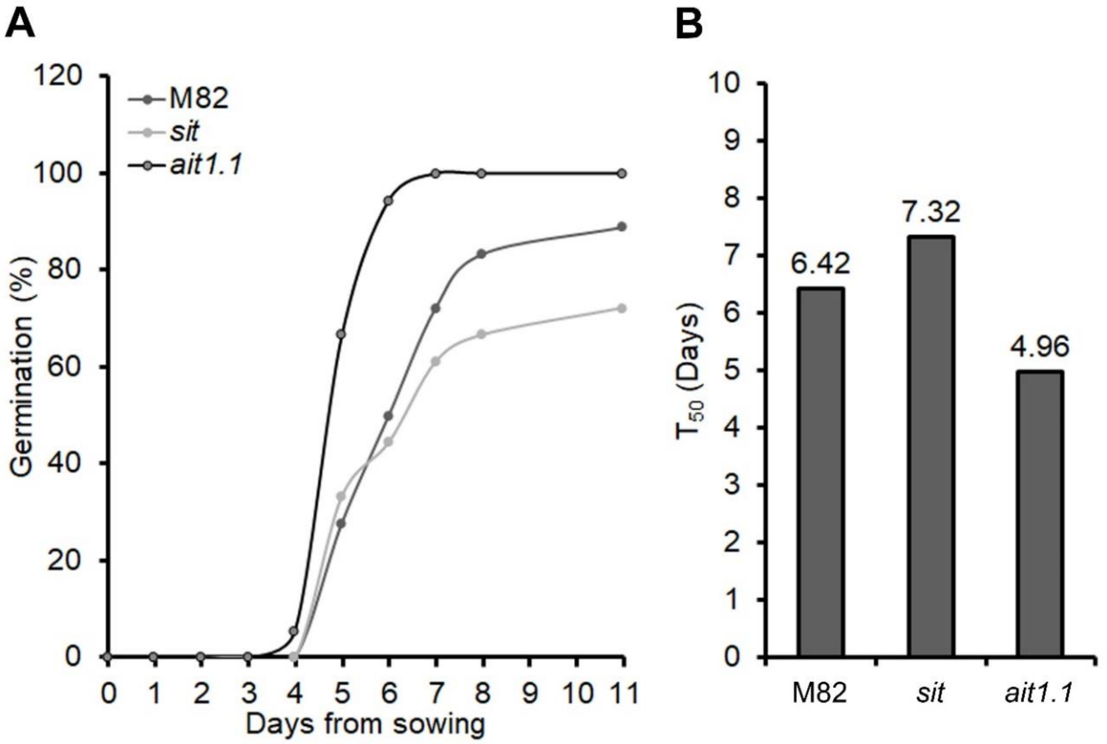


**Fig. S2** ABA deficiency reduced seed longevity. (A) Germination of M82, ABA-deficient mutant *sit* and *ait1.1* seeds stored for 18 months. (B) Germination rate ($T_{50}$) of M82, *sit* and *ait1.1*. Values are percentages (A) or means (B) of 50 seeds.

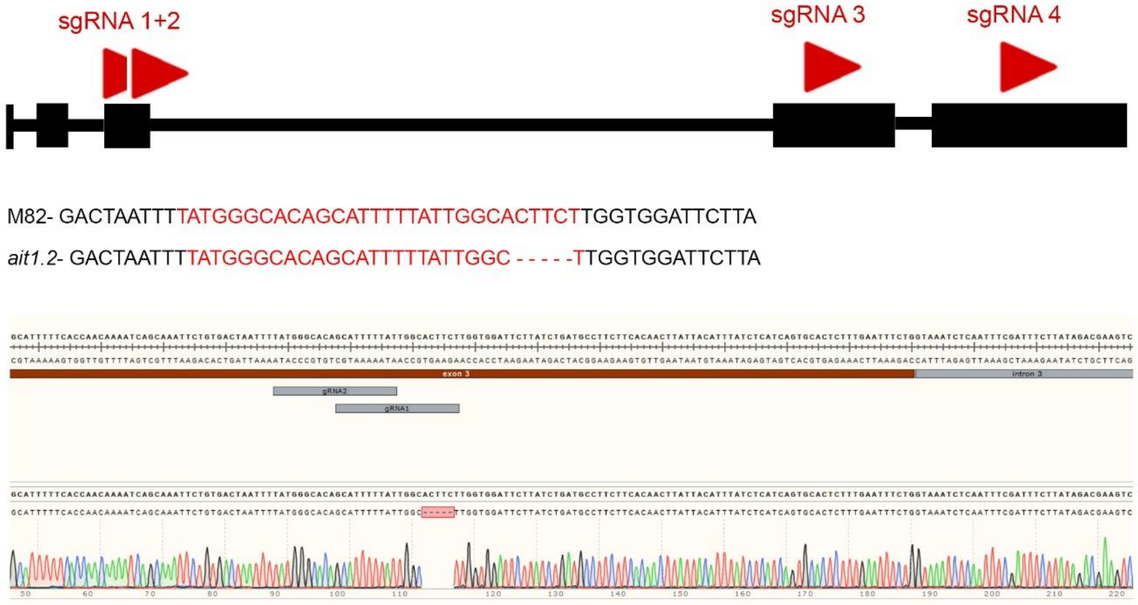


**Fig. S3** Sequence analysis of *ait1.2* CRISPR mutant allele*.* Schematic structure of *AIT1.2* gene with the positions of the RNA guides (red arrows) in the third, fourth and fifth exons. Also shown, the sequence of RNA guide1 and 2 (red letters), the chromatograms and the nucleotide sequences including the mutation that led to frame shift and premature stop codon.

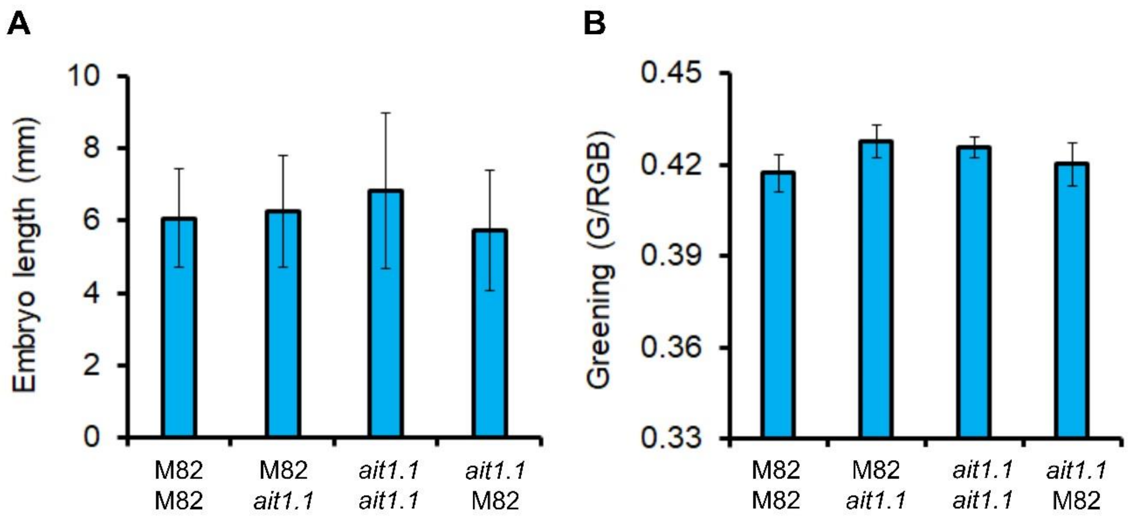


**Fig. S4** AIT1.1 did not affect endosperm inhibition of embryo growth. Length (A) and greening (B) of M82 and *ait1.1* isolated embryos, grown for 72h on a layer of endosperms. Values are means of 10 biological replicates ±SE.

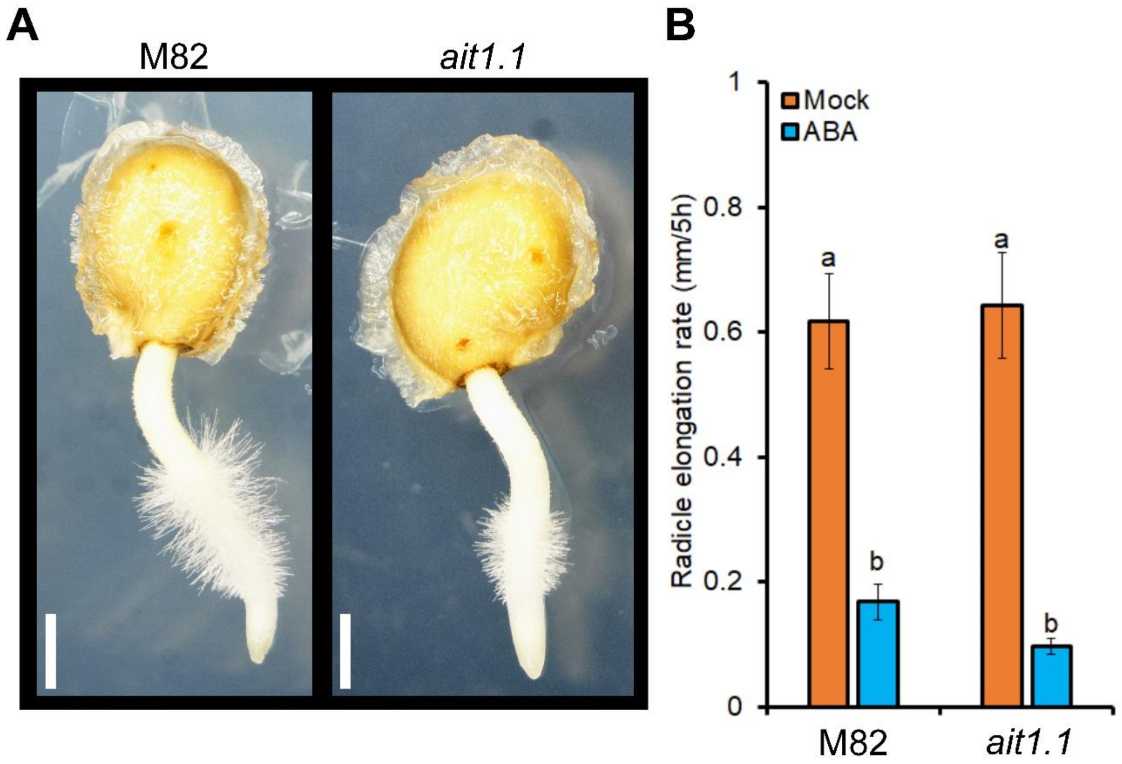


**Fig. S5** Loss of *AIT1.1* did not affect post-emergence radicle elongation. (A) Representative M82 and *ait1.1* post-emergence scarified seeds. (B) Elongation rate of post-emergence M82 and *ait1.1* radicles in scarified seeds, placed for 48h on MS medium. Values are means of 8 biological replicates ±SE. Small letters (B) represent significant differences between respective treatments by Tukey-Kramer HSD test ($P < 0.05$). Scale bar= 1 mm.

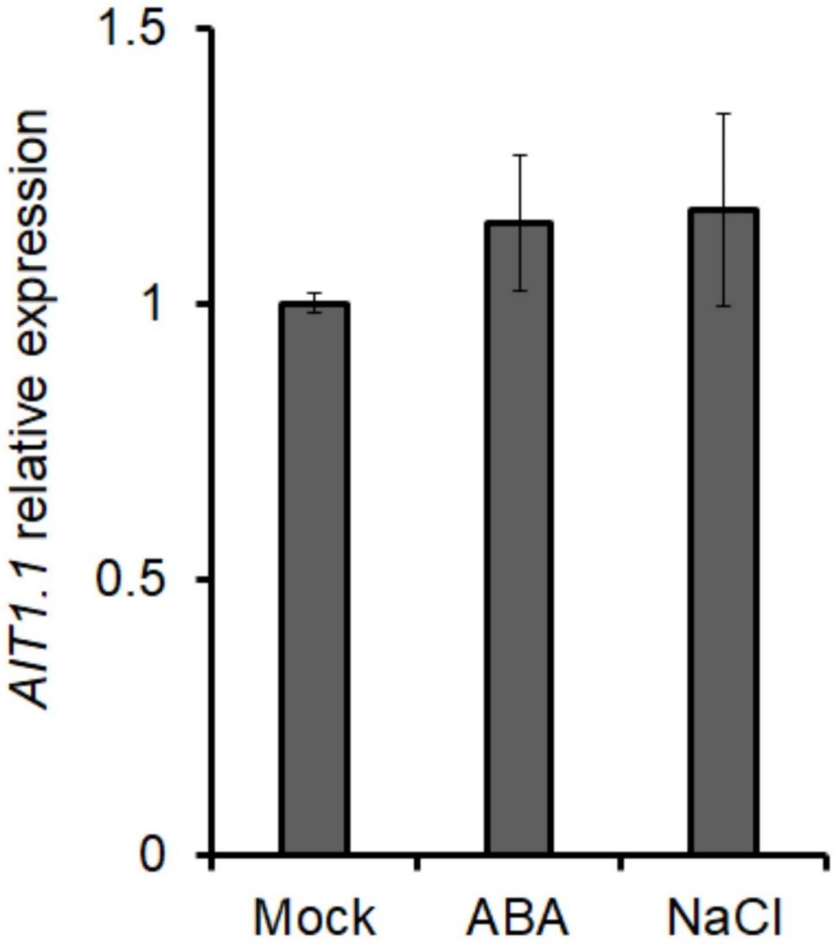


**Fig. S6** Salt or ABA treatments did not affect *AIT1.1* expression in imbibed seeds. Relative expression of *AIT1.1* in imbibed seeds treated for 48h with (or without) 50mM NaCl or 10µM ABA. Values are means of 4 replicates each contains 10 seeds ±SE.

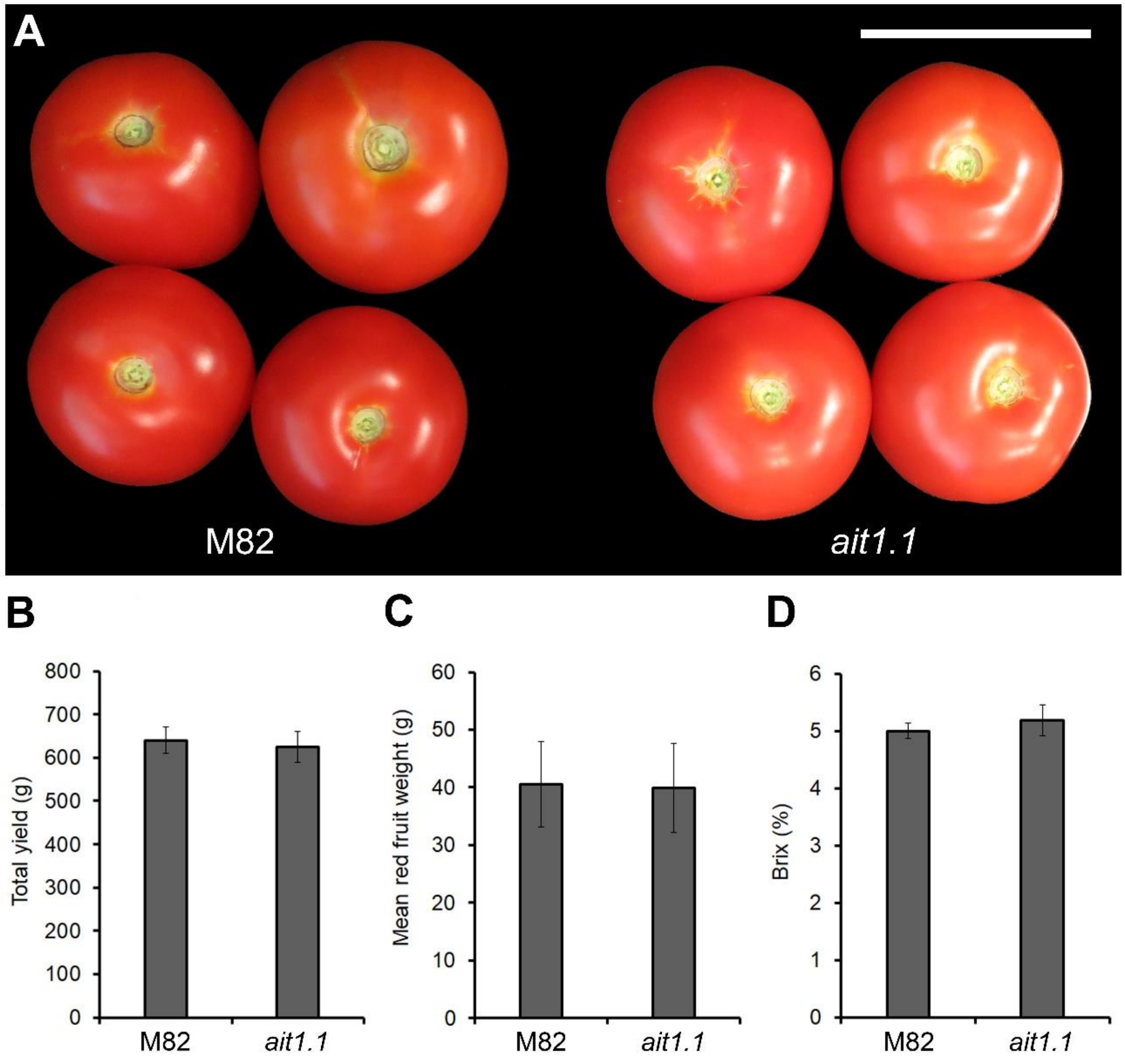


**Fig. S7** Loss of *AIT1.1* did not affect yield production. (A) Representative M82 and *ait1.1* red-ripen fruits. (B) Total fruit yield per plant (green and red fruits). (C) Mean weight of red-ripen fruit. (D) Brix of red-ripen fruits. Values are means of 20 plants ±SE. Scale bar in A = 5 cm.

**Supplemental Tables**

**Supplemental Table 1.** Primers used in this study.

| Gene | Used for | Sequence (5'-3') |
|---|---|---|
| *ACTIN* | RT-qPCR | Forward- GTCCTCTTCCAGCCATCCAT<br>Reverse- ACCACTGAGCACAATGTTACCG |
| *AIT1.1* | RT-qPCR | Forward- GTGGAAGCCCTCTCACAACA<br>Reverse- CTTCGCCATTTTCCTTGCCC |
| *AIT1.2* | RT-qPCR | Forward- AAGCAGCCTCGATGAACACA<br>Reverse- GGGGAAAACAGGGAGGGAAG |
| *AIT1.3* | RT-qPCR | Forward- GGCATTGGGTGTTGGAGGTA<br>Reverse- TCGCCGTCAAATTGTTCAGC |
| *AIT1.4* | RT-qPCR | Forward- CTTGTGGCGGCATTTTCCAA<br>Reverse- CCATGTCCTTCTTCCGCACT |
| *ABI3* | RT-qPCR | Forward- TTGGGTATGTTGGCCTTCTC<br>Reverse- TCTGCTGGTTCTGTGATTGC |
| *FUS3* | RT-qPCR | Forward- GCGGTATTCCGAGGTTATGA<br>Reverse- GTTGTCCATTGCAGGGAAGT |
| *DOG1* | RT-qPCR | Forward- CGAGCTAAACGACGACGAAT<br>Reverse- CACCAAGTAGGGGCAAAGAA |
| *LEC1* | RT-qPCR | Forward- GCTACCGTAGGTTCCCACAA<br>Reverse- TCGCGTCGTCTGATATCTTG |

| | | |
|---|---|---|
| *β1-3-glucanase* | RT-qPCR | Forward- AATGCAGCAACATGCTTGAG<br>Reverse- GGGGGAAAACAGACCAAAAT |
| *EXPA2* | RT-qPCR | Forward- GTGCAATGTCAATGCTGACC<br>Reverse- TTAGCCAGGGCATAGTTTGG |
| *MAN2* | RT-qPCR | Forward- AGGAACGATGGGAGGAAGTT<br>Reverse- CCAGCAGTGGATGGATTTTT |

**Supplemental Table 2.** gRNAs used in this study.

| Gene | Used for | Sequence (5'-3') |
|---|---|---|
| *AIT1.2* gRNA-1 | CRISPR | GCATTTTTATTGGCACTTCT |
| *AIT1.2* gRNA-2 | CRISPR | TATGGGCACAGCATTTTTAT |
| *AIT1.2* gRNA-3 | CRISPR | TTAGGCGTTGGAGGTATAAA |
| *AIT1.2* gRNA-4 | CRISPR | CGTAGTATTCATCATGATCT |
